# Catalpol inhibits HHcy-induced EndMT in endothelial cells by modulating ROS/NF-κB signaling

**DOI:** 10.1101/2023.06.05.543810

**Authors:** Chengyan Wu, Rusheng Zhao, Yuanhao Li, Peixia Li, Huibing Liu, Libo Wang, Xuehui Wang

**Affiliations:** Department of Cardiology, The First Affiliated Hospital of Xinxiang Medical University, Heart Center of Xinxiang Medical University, Xinxiang, China; College of Chemistry and Chemical Engineering, Henan Normal University, Xinxiang, China

**Keywords:** Hyperhomocysteinemia, endothelial dysfunction, vascular disease, endothelial mesenchymal transition, catalpol

## Abstract

**Background:** Hyperhomocysteinemia (HHcy) is an independent risk factor for atherosclerosis (AS), the molecular mechanisms of its pathogenesis are not fully understood. Endothelial dysfunction is the key initiating link in AS. However, whether endothelial-mesenchymal transition (EndMT) is involved in the regulation of HHcy-induced endothelial dysfunction and the role of catalpol in it remain unexplored.

**Methods and Results:** *In vitro* HHcy-treated primary human umbilical vein endothelial cells (HUVECs) were used to construct a model of endothelial dysfunction, and the antioxidants N-acetylcysteine (NAC) and catalase alcohol were administered. The endothelial dysfunction model was constructed by observing cell morphological changes, measuring intracellular reactive oxygen species (ROS) content, quantifying EndMT marker proteins (VE-cadherin, α-SMA, N-cadherin) and p-p65 and p65 protein expression by western blot, and detecting VE-cadherin and α-SMA fluorescence intensity by immunofluorescence. *In vivo* C57BL/6N mice were given a diet fed with 4.4% high methionine chow to construct a HHcy mice model and were treated with catalpol. The model was further validated by measuring the concentration of homocysteinemia (Hcy) in mice, HE and Masson staining to observe the pathological changes in the endothelium of aortic roots, immunohistochemistry to detect the expression of EndMT marker protein, immunofluorescence to detect the fluorescence intensity of p-p65, and fluorescent probe method to detect the content of ROS in the endothelium of aorta. The results showed that catalpol significantly inhibited HHcy-induced endothelial cell morphological transformation, reduced HHcy-induced increase in intracellular ROS content and α-SMA, N-cadherin, p-p65 protein expression, increased HHcy-induced decrease in VE-cadherin, CD31 protein expression, and was able to protect against endothelial pathological changes in the aortic root and reduce aortic endothelial ROS content.

**Conclusions:** Catalpol inhibits HHcy-induced EndMT, and the underlying mechanism may be related to the ROS/NF-κB signaling pathway. Catalpol may be a potential drug for the treatment of HHcy-related AS.

## 1 Introduction

With the increasing standard of living and aging of society, cardiovascular diseases, especially ischemic heart disease and stroke, have become the leading causes of death worldwide, accounting for 85% of the total number of deaths and healthcare costs worldwide.^1^ Coronary atherosclerotic heart disease is the most common ischemic heart disease. It is caused by narrowing or blockage of the lumen due to plaque in the vessel wall following impaired endothelial function or dysfunction of coronary arteries, resulting in myocardial ischemia, hypoxia, or necrosis. Hcy is a sulfur-containing, non-protein amino acid formed during methionine metabolism. Impaired Hcy metabolism and genetic or nutritional defects can lead to elevated plasma concentrations of Hcy and its formation. HHcy is considered an independent risk factor for AS and was first described by McCully (1969) in a study of two people who had been diagnosed with cystathionine beta Hcyuria.^2^ In addition, atherosclerotic plaques were discovered in two children with HHcy due to cystathione-β deficiency and defective vitamin B^12^ metabolism.^2^ This phenomenon has triggered extensive research on HHcy-induced vascular diseases. Subsequent studies verified that HHcy administered subcutaneously to New Zealand white rabbits for 5 weeks or fed a high methionine diet for 12 weeks resulted in the formation of AS plaques in the aorta of rabbits.^3,4^ In recent years, studies on HHcy-induced AS lesion formation have continued and the same conclusions have been reached.^5,6^ The following hypotheses have been suggested for the link between HHcy and endothelial dysfunction and AS: interference of HHcy with nitric oxide production;^7,8^ deregulation of the hydrogen sulfide signaling pathway;^9,10^ oxidative stress, inflammation, and impaired lipoprotein metabolism;^11,12^ protein N-homocysteinylation;^13,14^ and cellular hypomethylation^15,16^, among others.^17^

Atherosclerotic plaques are formed by the accumulation of lipids, mesenchymal cells, immune cells, and the extracellular matrix, among which the accumulation of mesenchymal cells (including myofibroblasts, fibroblasts, fibroblasts, and smooth muscle cells) is an important component of plaque formation. The origin of plaque mesenchymal cells has been extensively studied for many years. Evrard used an endothelial-specific lineage tracking system and found that a large number of plaque-associated mesenchymal cells were derived from endothelial cells.^18^ Endothelial cells undergo a series of molecular events to lose endothelial cell morphology and characteristic gene expression, and gain phenotypic characteristics. Gene expression associated with mesenchymal cells is termed EndMT.^19^ It is a specific form of epithelial–mesenchymal transition (EMT). In recent years, there has been increasing evidence that EndMT is involved in the development of cardiovascular diseases, including AS, pulmonary hypertension, valve disease, remodeling after vascular injury, and myocardial fibrosis.^20^ EndMT activation is believed to involve several signaling pathways, including TGF-β signaling, oxidative stress and inflammation, cellular metabolism, non-coding RNA, epigenetics, Wnt/β-linked protein signaling, Notch signaling, and fibroblast growth factor.^21^

Recent studies have reported that EndMT may be involved in regulating HHcy-mediated endothelial dysfunction, and that this process can be inhibited by the Chinese patented medicines ganoderma lucidum triterpenes and rhodiol glycosides.^22,23^ However, this study has not been conducted in animal experiments or clinical settings. In the renal system, Li et al. observed that glomerular podocytes in mice with HHcy model showed EMT, which was dependent on the NOX/HIF-1α signaling pathway and could be inhibited by growth hormone treatment.^24^ In addition, Zhang et al. performed *in vivo* experiments to validate the hypothesis that HHcy induces EMT in glomerular podocytes through activation of NADPH oxidase.^25^ Li et al. reported similar results in *in vitro* experiments.^26^ These studies provided an experimental basis for the involvement of EndMT in the regulation of HHcy-induced endothelial dysfunction.

Catalpol, an active component of groundnuts, has the highest content in fresh groundnuts and belongs to the group of cyclic enol ether terpene glucosides with the molecular formula C_15_H_22_O_10_, as shown in (Figure 7a). Several studies have shown that catalpol has therapeutic, cardiovascular-protective, neuroprotective, anticancer, and hepatoprotective effects in the treatment of diabetes and that its biological functions are mainly related to its anti-inflammatory and antioxidant effects.^27^ In the cardiovascular system, catalpol exerts its anti-atherosclerotic effects mainly through the inhibition of oxidative stress, inflammatory response, neoplastic endothelial proliferation, macrophage infiltration, lipid metabolism, anti-fibrosis, and the reduction of extracellular matrix aggregation.^28^

Based on the above background, the specific molecular mechanism of HHcy-induced vascular dysfunction is not fully understood, and it is not known whether catalpol exerts a protective effect on it. The objectives of this study were: 1) to investigate whether HHcy induces EndMT *in vitro* and *in vivo*; 2) to investigate the role of catalpol in HHcy-induced EndMT; and 3) to investigate whether catalpol regulates HHcy-induced EndMT through the ROS/NF-κB signaling pathway.

## 2 Materials and methods

### 2.1 Materials

Hcy was purchased from Sigma-Aldrich (item no. H4628, ready-to-use); catalpol was purchased from Jingzhu Biological (item no. Z100599-1g) and diluted with autoclaved pure water to a concentration of 3 mM master mix, packed in eppendorf (EP) tubes, and frozen at −80°C for 1 week; primary HUVECs were purchased from Science Cell, USA (item no. DFSC-EC-01). ECM complete medium was purchased from Science Cell, USA (item no. 1001) and 4.4% high methionine feed was purchased from Jiangsu Synergy Pharmaceutical and Biological Engineering Co.

Anti-GAPDH, VE-cadherin, N-cadherin, α-SMA (smooth muscle actin), and CD31 antibodies were purchased from Proteintech. Horseradish peroxidase (HRP)-coupled goat anti-rabbit IgG was purchased from Jackson ImmunoResearch Laboratories. p-p65 protein antibodies were purchased from Cell Signaling Technology and p65 protein antibodies were purchased from Wan Lei Biology. NAC was purchased from Selleck (item no. 1623). Cellular ROS assay kits, BCA protein quantification kits, and fluorescence-labeled secondary antibodies were purchased from Shanghai Biyuntian Biological Co. The CCK8 kit was purchased from Abbkine. The reactive oxygen detection kit for tissue sections was purchased from Shanghai Pebble Biotechnology Company. Hematoxylin & eosin (HE) stain and modified Masson stain were purchased from Beijing Solaibao Technology Co.

### 2.2 Cell experiments and methods

#### 2.2.1 Cell culture

Cells were grown in ECM complete medium (containing 5% FBS, 1,000 μg/ml P/S, 1% ECGS) and cultured in a 5% CO_2_ incubator. When the cell density grew to ∼70%–80%, the cells were digested and passaged using trypsin; cells were passaged 2–3 times before the experiment. Control group cells were cultured by adding ECM complete culture medium. Hcy group cells were cultured by adding cell culture medium pre-mixed with certain concentrations of Hcy. Treatment group cells were co-treated with Hcy and Catalpol. Hcy+NAC group cells were co-treated with Hcy and 5 mM NAC.

#### 2.2.2 Cell viability assay

Cells were inoculated in 96-well plates, HUVECs were treated with different concentrations of Hcy (0, 50, 100, 200, 400, and 800 μM), and cell viability was measured using the CCK-8 reagent. We established 5–7 replicate wells in each group, and each well was incubated in an incubator at 37℃ for 1–4 h after the addition of 10 ul CCK8 reagent. The absorbance value of each well at 450 nm was measured using an enzyme standardization instrument, and the experiment was repeated at least three times.

#### 2.2.3 Cell morphology observation

Cells were inoculated in 6-well plates at approximately 6,000 cells/well, and cell morphological changes were observed after co-treatment with or without Hcy (800 μM), Hcy+Catalpol (30 μM), and Hcy+NAC (5 mM), and observed under a microscope and image acquisition.

#### 2.2.4 Western blot analysis

Cells were inoculated in 6-well plates and rinsed twice with PBS after drug treatment. The protein lysis solution was added on ice, scraped, and transferred to EP tubes, lysed three times using an ultrasonic lyser, and proteins were denatured by heating at 100°C for 10 min, naturally cooled to room temperature, centrifuged at 4°C and 12,000 rpm/min for 5 min, and set aside in a −80°C refrigerator. The protein concentration was determined using the bicinchoninic acid method. After electrophoresis, the membranes were transferred using the wet method and incubated with 5% skim milk for 1 h. The membranes were incubated overnight with the corresponding antibodies (VE-cadherin 1:2,000, N-cadherin 1:2,000, α-SMA 1:1,000, p-p65 1:1,000, and p65 1:1,000) at a certain ratio. The membranes were washed three times with TBST, incubated with horseradish peroxidase-coupled secondary antibody (1:2,000) for 1–1.5 h, washed again three times with TBST, and configured for color development solution exposure. The ImageJ software was used to analyze the grayscale values. The corresponding target protein was expressed as the ratio of the gray value of the target protein band to the gray value of the internal reference (GAPDH).

#### 2.2.5 Immunofluorescence staining

The cells were inoculated into 24-well plates, incubated for 6 h to make the cells adhere to the wall, and treated with the added drugs (control, Hcy, Hcy+Catalpol groups) for 48 h. The cells were fixed with 4% paraformaldehyde for 10 min, washed three times with PBS, permeabilized with 0.1% Triton X-100 for 10 min, washed three times with PBS, and blocked with 5% bovine serum albumin (BSA) for 30 min. The primary antibody (VE-cadherin 1:500, α-SMA 1:800) in a certain ratio was added dropwise and incubated overnight at 4℃. It was then washed three times with PBST, fluorescent secondary antibody (1:500) was added dropwise for 1 h (after this operation, light was avoided), washed three times with PBST, stained with DAPI (1 ug/ml) for 15 min, washed three times with PBS, and a slice was sealed with anti-quenching sealer. Finally, the films were observed under a laser confocal microscope or fluorescence microscope, and image acquisition and fluorescence intensity analysis were performed using ImageJ.

#### 2.2.6 Oxidative stress assessment

The DHE master mix (10 mM) was diluted using serum-free ECM medium at a dilution ratio of 1:1,000 to prepare an ROS staining working solution. The treated six-well plate cells were washed two times using PBS; then, 1 ml/well of DHE working solution was added, and incubation was continued for 20–30 min at 37℃ before washing three times using PBS. Finally, the cells were observed under a fluorescence microscope and images were acquired; the fluorescence intensity was analyzed using ImageJ.

### 2.3 Animal experiments and methods

#### 2.3.1 Animal model construction and sample treatment

We fed 18 healthy male C57BL/6N mice of 6–8 weeks old with water ad libitum at room temperature 22℃ ± 1℃ with appropriate humidity. After 1 week of acclimatization, the mice were randomly divided into three groups at week 2: control group (general diet), high methionine (Met) group (4.4% methionine diet), and treatment group (4.4% Met diet + Catalpol 20 mg/kg/d), with six mice in each group. Control mice were fed a normal diet ad libitum; model mice were given a 4.4% high-Met diet ad libitum; treated mice were given a 4.4% high-Met diet ad libitum + catalpol dilution 20 mg/kg/d intraperitoneally. The mice were co-fed for 14 weeks. At the end of feeding, mice were anesthetized with 3% sodium pentobarbital (40 mg/kg); 1–1.5 ml of blood was taken from the eye of each mouse, left at room temperature for ∼30 min, then centrifuged at 3,000 rpm/min for 15 min to separate the upper layer of serum, which was cryopreserved and sent to Zhengzhou Dean Medical Laboratory for detection of serum Hcy concentration. At the same time, the heart and aorta of mice were quickly removed and flushed with pre-cooled PBS. The heart was fixed in 4% paraformaldehyde and used for HE staining, modified Masson staining, and immunohistochemistry and immunofluorescence analysis. The aorta was temporarily stored in 0.9% saline; immediately frozen sections of tissue were used for ROS detection.

#### 2.3.2 HE staining

The fixed heart tissues were routinely paraffin embedded, sliced to 4–5 μm thickness, set aside, and then baked for 1–1.5 h. Xylene (I) for 10 min→xylene (II) for 10 min→100% ethanol for 3 min→95% ethanol for 3 min→85% ethanol for 3 min→75% ethanol for 3 min→50% ethanol for 3 min→distilled water soaking for 2 min. Samples were then stained in hematoxylin staining solution for 2 min, washed with distilled water to remove floating color, subjected to differentiation solution for 10–60 s, rinsed with tap water three times, stained with eosin staining solution for 40 s, washed with tap water to remove staining solution. The slices were routinely dehydrated, transparent, and sealed with neutral gum. Samples were observed under a light microscope and images were acquired.

#### 2.3.3 Modified Masson staining

The fixed heart tissues were routinely paraffin-embedded, sectioned to 4–5 μm thickness, baked for 1–1.5 h, and routinely dewaxed to water. Coal staining solution was drip-stained overnight at room temperature; azurite staining solution was drip-stained for 2–3 min and washed twice for 10–15 s each time. Mayer hematoxylin staining solution was drip-stained for 2–3 min and washed twice for 10–15 s each time. Acidic differentiation solution was used to differentiate for a few seconds and then washed away to terminate differentiation; then, the samples was rinsed in distilled water for 10 min. Lichon red magenta staining solution was drip-stained for 10 min and washed twice for 10–15 s each time The slices were treated with phosphomolybdic acid solution for ∼10 min; then, the upper solution was poured off, and the sections were directly stained with aniline blue staining solution for 5 min without washing. After the aniline blue solution was washed off with weak acid, the sections continued to be covered with weak acid working solution dropwise for 2 min; the sections were routinely dehydrated, transparent, and sealed with neutral gum. The sections were observed under an optical microscope, images were acquired, and collagen fiber deposition was analyzed using ImageJ.

#### 2.3.4 Immunohistochemistry staining

The fixed heart tissues were routinely paraffin-embedded, sectioned to 4–5 μm thickness, baked for 1–1.5 h, and routinely dewaxed to water. The sections were immersed in citrate buffer for 10 min, heated in a microwave for antigen repair, naturally cooled to room temperature, and washed three times with PBS. The sections were immersed in 0.2% Triton X-100 for 10 min for permeabilization, and washed three times with PBS. Next, they were immersed in 3% H_2_O_2_ for 10 min to inactivate peroxidase, and washed three times with PBS. Then, they were closed in 10% BSA serum for 1 h. Primary antibody (CD31 1:1,000, VE-cadherin 1:800, α-SMA 1:1,500), configured according to a certain ratio, was added dropwise and the sections were incubated overnight at 4℃ before washing three times with PBS. Next, they were treated with dropwise addition of secondary antibody and incubated for 1 h in a wet box at room temperature before washing three times with PBS. Next, they were treated with dropwise addition of DAB chromogenic solution for color development (time varied from seconds to 10 min). Hematoxylin re-staining was then performed for 40 s. The slices were routinely dehydrated, made transparent, and sealed with neutral gum. Images were acquired under a light microscope for the quantitative analysis of target proteins using ImageJ.

#### 2.3.5 Immunofluorescence staining

The fixed heart tissues were routinely paraffin-embedded, sectioned to 4–5 μm thickness, baked for 1–1.5 h, and routinely dewaxed to water. The sections were then immersed in citrate buffer for 10 min, heated in a microwave for antigen repair, cooled naturally to room temperature, and washed three times with PBS. They were then soaked in 0.2% Triton X-100 for 10 min for permeabilization before washing three times with PBS. Next, they were closed with 10% BSA serum for 1 h. The primary antibody (p-p65 1:800) was added dropwise and sections were incubated overnight at 4ºC before washing with PBS three times. Fluorescent secondary antibody (1:500) was added dropwise and the sections were incubated for 1 h at room temperature in a wet box before washing three times with PBS. DAPI was added dropwise and incubated for 15 min at room temperature before washing two times with PBS. The slices were then sealed with anti-fluorescence quenching agents. Images were obtained using a laser confocal microscope or fluorescence microscope; ImageJ was used for fluorescence intensity analysis.

#### 2.3.6 ROS detection in frozen tissues sections

Dissected and separated fresh aorta was cut into small segments of ∼1 cm, placed flat in a special cassette (diameter ∼2 cm) in the frozen section machine, submerged by adding an appropriate amount of OCT embedding agent, and then cut into sections of 10–20 μm thickness in a frozen section machine after complete solidification. At room temperature, a 200 μl/slice of washing solution was added dropwise to the sections and left for 3–5 min; the washing solution was carefully aspirated and a 100–200 μl/slice of staining probe solution was added dropwise and incubated at 37°C for 20–60 min in an incubator protected from light. Next, the staining solution was removed, and the sections were washed three times with PBS and then sealed with glycerol. The sections were observed under a laser confocal microscope or fluorescence microscope,; images were acquired and fluorescence intensity was analyzed using ImageJ.

### 2.4 Statistical analysis

GraphPad Prism 8.0.2 was used for statistical analysis and the creation of statistical plots. Normally distributed measures were expressed as mean ± standard deviation (SD), and comparison of means between multiple groups was performed by one-way analysis of variance (ANOVA) with Tukey’s multiple comparisons. Differences were considered statistically significant at P < 0.05.

### 3 Results

### 3.1 HHcy induces EndMT in endothelial cells *in vitro*

To assess the effect of Hcy on cell viability in *in vitro* experiments, we treated HUVECs with 0, 50, 100, 200, 400, and 800 μM of Hcy, respectively. There were no significant differences in cell viability between groups (Figure 1a). To verify whether Hcy could induce EndMT in endothelial cells *in vitro*, we performed an EndMT-related protein expression assay 48 h after Hcy treatment. The results showed that VE-cadherin protein expression gradually decreased, and N-cadherin and α-SMA protein expression gradually increased with increasing Hcy concentration in a concentration-dependent manner (Figure 1b, 1c). A Hcy concentration of 800 μM was selected for subsequent experiments. To investigate the effect of Hcy on cell morphology, morphological changes in cells in the blank and Hcy groups were observed at 24, 48, and 72 h after treatment. The results showed that There was no significant difference in the percentages of mesenchymal cells (spindle cells) between the two groups after 24 h; however, the number of spindle cells was significantly higher in the Hcy group than in the blank group after 48 and 72 h (Figure 1d). This indicates that HUVECs shifted from the cobblestone morphology of endothelial cells to the spindle-shaped morphology of mesenchymal cells upon Hcy induction. These data suggested that HHcy induces EndMT in endothelial cells *in vitro*.

**Figure 1.**
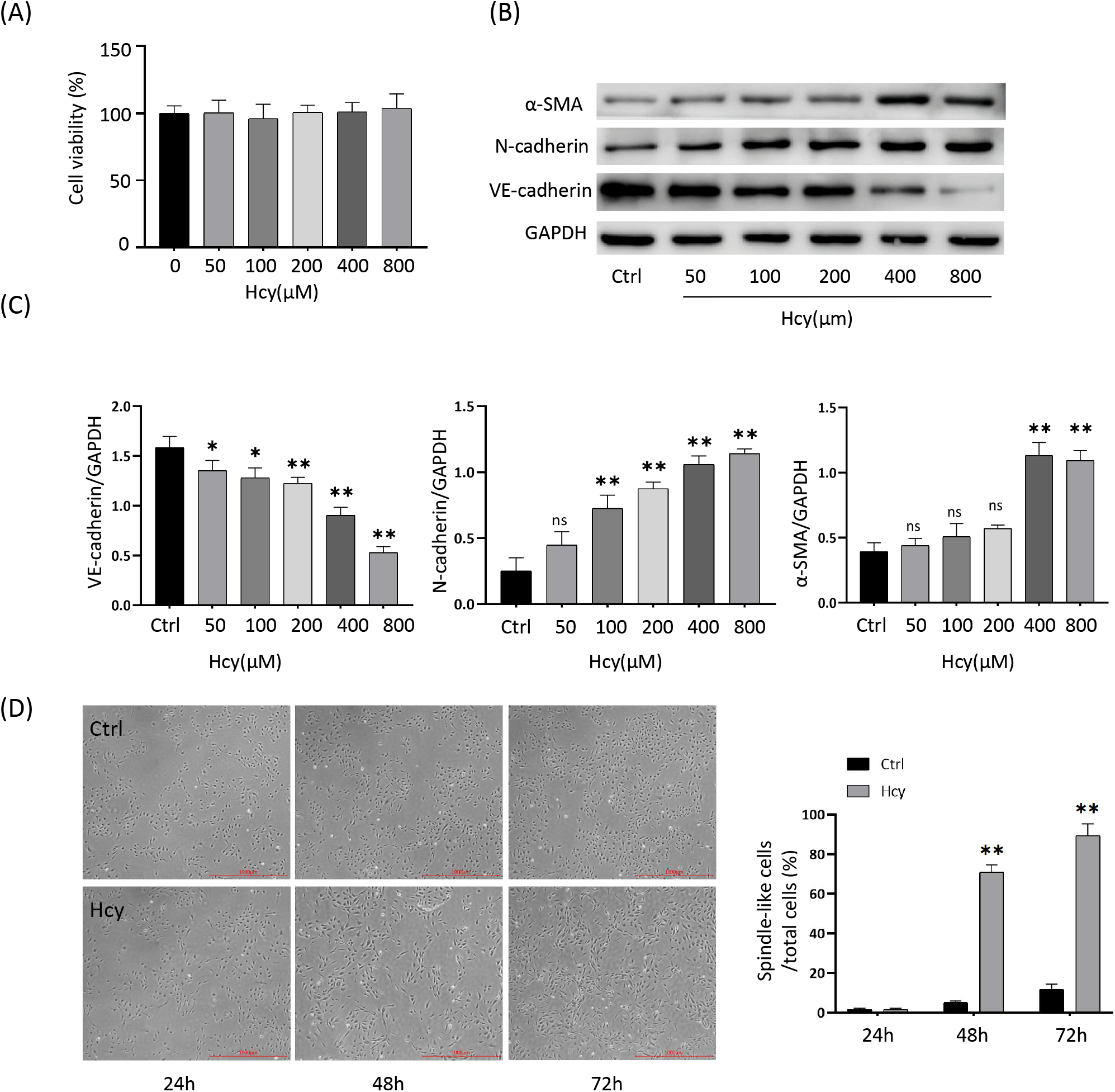
HHcy induces EndMT in endothelial cells *in vitro*. (**A**) Cell viability after treatment of HUVECs with different concentrations of Hcy (0–800 umol/L). (**B**) Protein immunoblotting experiments for EndMT-related protein expression levels of VE-cadherin, N-cadherin, and α-SMA, and quantification of protein levels (lower panel) after treatment of HUVECs with different concentrations of Hcy (0–800 umol/L) for 48 h. (**C**) Cell morphology of HUVECs after treatment with or without Hcy (800 umol/L) for 24, 48, or 72 h, and the percentage of spindle cells to the total cell number (right panel). Data are expressed as mean ± SD, n = 3, *P < 0.05, ** P < 0.01, compared with the control group.

### 3.2 Catalpol inhibits HHcy-induced EndMT in endothelial cells

To investigate the effect of Catalpol on HHcy-induced EndMT in endothelial cells *in vitro*, we chose Catalpol at a concentration of 30 μM as the treatment dose. First, the effects of Hcy and Hcy+Catalpol groups on cell morphological changes were observed. The results showed that the number of spindle cells was significantly increased in the Hcy group compared with the blank group, and the number of spindle cells was significantly decreased in the Hcy+Catalpol group compared with the Hcy group (Figure 2a), indicating that HUVECs shifted from endothelial cell cobblestone morphology to mesenchymal cell spindle morphology under the induction of Hcy, and Catalpol could inhibit this cell morphology change to some extent. To further verify the role of Catalpol on Hcy-induced EndMT in endothelial cells *in vitro*, we performed protein immunoblotting and immunofluorescence staining experiments on cells 48 h after drug treatment to detect EndMT-related protein expression. The results showed that Hcy+Catalpol treatment increased the Hcy-induced decrease in VE-cadherin protein expression (fluorescence intensity), decreased the Hcy-induced increase in α-SMA protein expression (fluorescence intensity), and decreased the Hcy-induced increase in N-cadherin protein expression (Figure 2b, 2c). These results suggest that catalpol inhibits HHcy-induced EndMT in endothelial cells *in vitro*.

**Figure 2.**
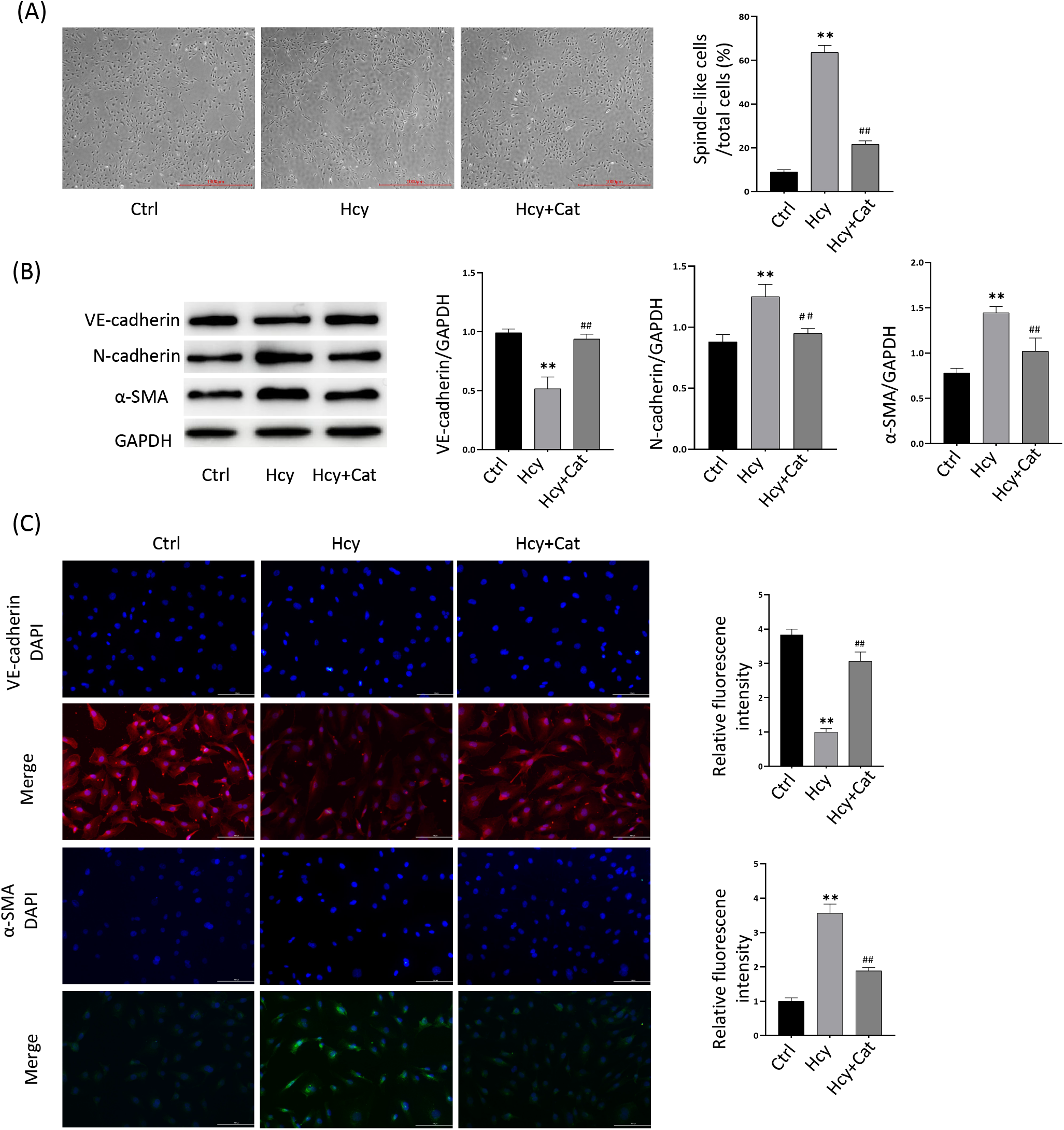
Catalpol inhibition HHcy induced EndMT in endothelial cells. (**A**) Cell morphology of Ctrl, Hcy, and Hcy+Cat groups after treatment of HUVECs for 48 h, and the percentage of shuttle cells to the total cell number (right panel). (**B**) Protein immunoblotting experiments showing expression levels of EndMT-related proteins VE-cadherin, N-cadherin, and α-SMA after 48 h treatment of HUVECs in the Ctrl, Hcy, and Hcy+Cat groups, respectively, and quantification of protein levels (right panel). (**C)** Immunofluorescence experiments showing VE-cadherin, and α-SMA protein fluorescence intensity after 48 h treatment of HUVECs in the Ctrl, Hcy, and Hcy+Cat groups, respectively, and relative fluorescence intensity quantification (right panel). Data are expressed as mean ± SD, n = 3, *P < 0.05, ** P < 0.01, compared with the control group. #P < 0.05, ##P < 0.01, compared with the model group.

### 3.3 Catalpol attenuates HHcy-induced EndMT in endothelial cells by regulating ROS/NF-κB signaling

Previous studies have shown that in addition to the classical TGF-β-dependent signaling involved in EndMT activation, non-TGF-β-dependent signaling pathways such as, oxidative stress, inflammatory signaling, cellular metabolism, non-coding RNA, epigenetics, Wnt/β-linked protein signaling, Notch signaling, and fibroblast growth factor, among others, are involved in activation (Kovacic et al., 2018). In that study, we focused on investigating whether catalase acts on HHcy-induced EndMT by regulating the oxidative stress-inflammatory signaling pathway (i.e., ROS/NF-κB). We used the well-known antioxidant NAC as a ROS inhibitor to verify the involvement of oxidative stress in EndMT. First, we observed the morphological changes in HUVECs after Hcy and Hcy + NAC (5 mM) treatment for 48 h. The results showed that the number of spindle cells was significantly increased in the Hcy group compared with the blank group; and the number of spindle cells was significantly decreased in the Hcy+NAC group compared with the Hcy group (Figure 3a). This indicates that HUVECs shifted from endothelial cell cobblestone morphology to mesenchymal cell spindle morphology under Hcy induction, and NAC could inhibit this cell morphology change to some extent. Second, EndMT-related proteins were detected in Hcy-and Hcy+NAC-treated cells at 48 h post-treatment. The results showed that the Hcy+NAC group increased the decrease of VE-cadherin protein expression induced by the Hcy group and decreased the increase of α-SMA and N-cadherin protein expression induced by the Hcy group (Figure 3b). To investigate whether catalpol acts on HHcy-induced EndMT by regulating ROS/NF-κB signaling, first, we detected intracellular ROS content after drug treatment using the DHE probe assay. The results showed that the intracellular ROS content was significantly higher in the Hcy group than in the blank group and significantly lower in the Hcy+Catalpol group than in the Hcy group (Figure 3c). p-p65 and p65 protein expression was detected in the cells after 12 h of Hcy and Hcy+Catalpol treatment. The results showed that the increase in p-p65 protein expression induced by Hcy was significantly reduced in the Hcy + Catalpol group, but there was no significant effect on p65 protein expression (Figure 3d). These results suggest that catalpol attenuates HHcy-induced EndMT in endothelial cells by inhibiting ROS/NF-κB signaling *in vitro*.

**Figure 3.**
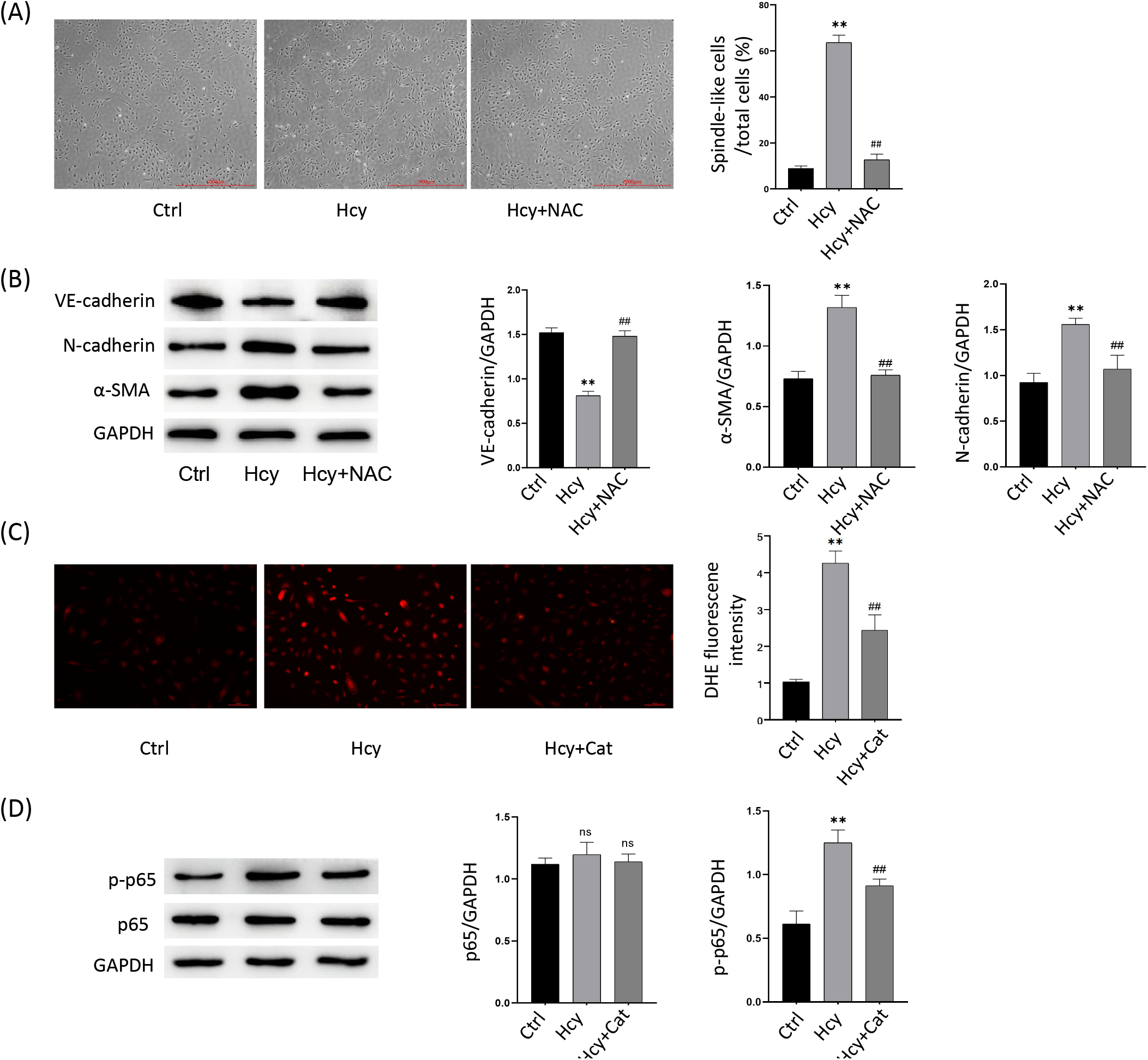
Catalpol attenuates HHcy-induced EndMT in endothelial cells by regulating ROS/NF-κB signaling. (**A**) Cell morphology of Ctrl, Hcy, and Hcy+NAC (5 mmol/L) after 48 h treatment of HUVECs in three groups, and the percentage of spindle-shaped cells to the total cell number (right panel). (**B**) Protein immunoblotting experiments showing expression levels of EndMT-related proteins VE-cadherin, N-cadherin, and α-SMA after 48 h treatment of HUVECs in Ctrl, Hcy, and Hcy+NAC groups, and quantification of protein levels (right panel). (**C**) DHE probe assay for intracellular ROS showing intracellular ROS levels after 48 h treatment of HUVECs in the Ctrl, Hcy, and Hcy+Cat groups, and quantification of fluorescence intensity (right panel). (**D**) Protein immunoblotting assay showing p-p65 and p65 protein expression levels after 12 h treatment of HUVECs in the Ctrl, Hcy and Hcy+Cat groups, and quantification of protein levels (right panel). Data are expressed as mean ± SD, n = 3, *P < 0.05, ** P < 0.01, compared with the control group. #P < 0.05, ##P < 0.01, compared with the model group.

### 3.4 Catalpol attenuates high Met-induced vascular endothelial dysfunction in mice *in vivo*

To investigate the effect of catalpol on high-Met-induced vascular endothelial dysfunction in mice *in vivo*, we constructed an HHcy model in C57BL/6N mice fed a 4.4% high-Met diet and treated them with or without catalpol 20 mg/kg/d intraperitoneally. The mice were anesthetized, sacrificed after 14 weeks of co-feeding, and sent to a third-party testing facility to detect serum Hcy concentrations, as shown in (Figure 4a). The results showed that feeding a high-Met diet significantly increased the serum Hcy concentration in mice and induced HHcy, but Catalpol treatment did not significantly reduce the Hcy concentration. This suggests that catalpol did not reduce Hcy levels by participating in the circulating metabolism of Hcy. Next, HE and modified Masson staining were performed on the isolated and fixed heart tissues to analyze morphological changes in the vascular endothelium and fibrosis. The results showed that the vascular endothelium of mice in the high-Met group showed a pyknotic morphology and fibrotic manifestations compared with the control group, whereas catalpol treatment significantly attenuated the high-Met-induced endothelial pathological changes in mice (Figure 4b, 4c). These data suggest that catalpol exerts a protective effect against high Met-induced vascular endothelial dysfunction in mice.

**Figure 4.**
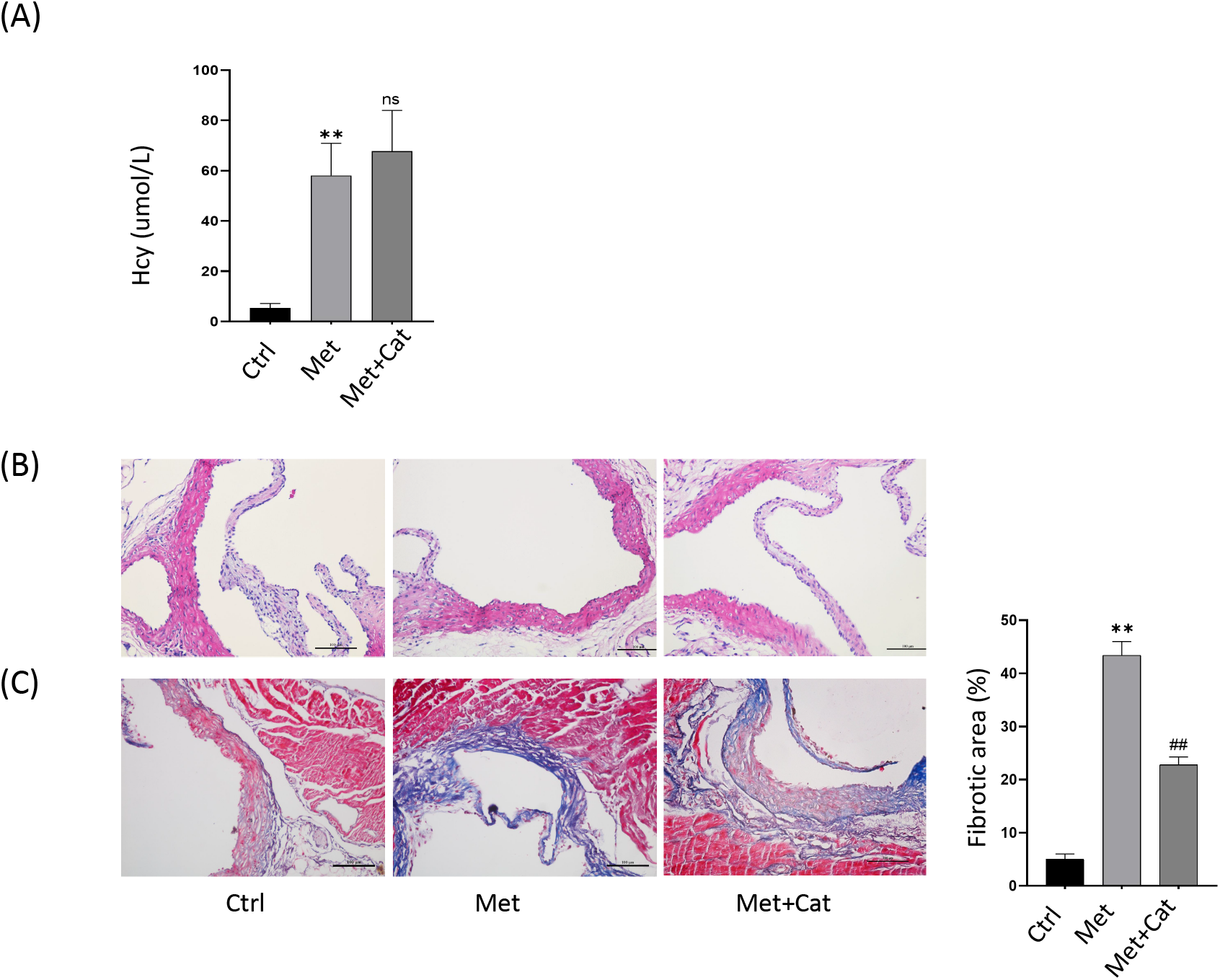
Catalpol attenuates high Met-induced vascular endothelial dysfunction in mice *in vivo*. (**A**) Serum Hcy concentration levels of mice in the three groups. (**B**) HE staining showing the pathomorphological changes in the endothelium of the aortic root in the three groups of mice. (**C**) Masson staining showing the endothelial collagen fiber deposition in the aortic root of the three groups of mice, and the percentage of collagen fiber area (right panel). Data are expressed as mean ± SD, n = 3, *P < 0.05, ** P < 0.01, compared with the control group. #P < 0.05, ##P < 0.01, compared with the model group.

### 3.5 Catalpol attenuates high Met-induced vascular endothelial EndMT in mice

To investigate the role of catalpol in high Met-induced endothelial dysfunction in mice *in vivo*, immunohistochemical staining of isolated and fixed heart tissues was performed to analyze changes in endothelial EndMT-related protein expression. The results showed that endothelial CD31 and VE-cadherin protein expression decreased and α-SMA protein expression increased in the high-Met group compared with the control group, whereas endothelial CD31 and VE-cadherin protein expression increased and α-SMA protein expression decreased in the catalpol-treated group compared with the high-Met group (Figure 5). These data suggest that high-Met feeding can induce the disruption or loss of vascular endothelial continuity and the EndMT process in mice, and that catalpol can inhibit this process to some extent and exert a protective effect.

**Figure 5.**
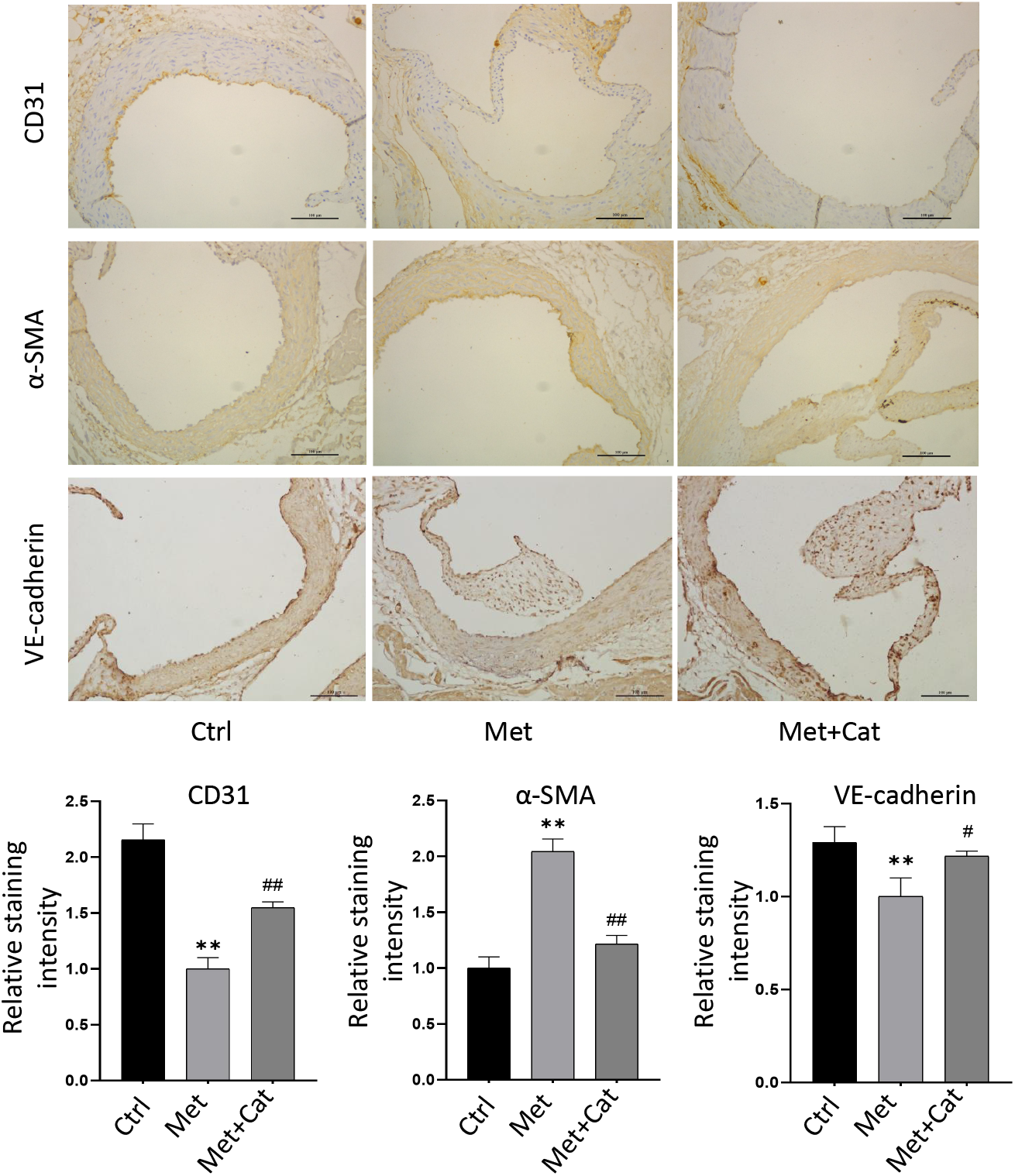
Catalpol attenuates high Met-induced vascular endothelial EndMT in mice. Immunohistochemical staining of EndMT-related protein expression of CD31, VE-cadherin, and α-SMA in the endothelium of mouse aortic roots in the control, high-Met, and high-Met + Catalpol groups, and quantification of protein (lower panel). Data are expressed as mean ± SD, n = 3, *P < 0.05, ** P < 0.01, compared with the control group. #P < 0.05, ##P < 0.01, compared with the model group.

### 3.6 Catalpol attenuates high Met-induced EndMT by modulating ROS/NF-κB signaling

In order to investigate whether catalpol plays a role in high Met-induced endothelial EndMT in mice by modulating ROS/NF-κB signaling, tissue reactive oxygen species staining and immunofluorescence staining were performed on isolated and fixed heart tissues to analyze the endothelial ROS content and p-p65 protein expression changes. The results showed that the vascular endothelial ROS content was significantly higher in the high-Met group than in the control group, and the catalpol-treated group had significantly lower vascular endothelial ROS content than the high-Met group (Figure 6a). In addition, immunofluorescence staining results showed that vascular endothelial p-p65 fluorescence intensity increased in the high-Met group and decreased in the catalpol-treated group compared with the high-Met group (Figure 6b). These data suggest that catalpol attenuates high Met-induced endothelial EndMT in mice by inhibiting ROS/NF-κB signaling. This is consistent with the results obtained from the *in vitro* experiments.

**Figure 6.**
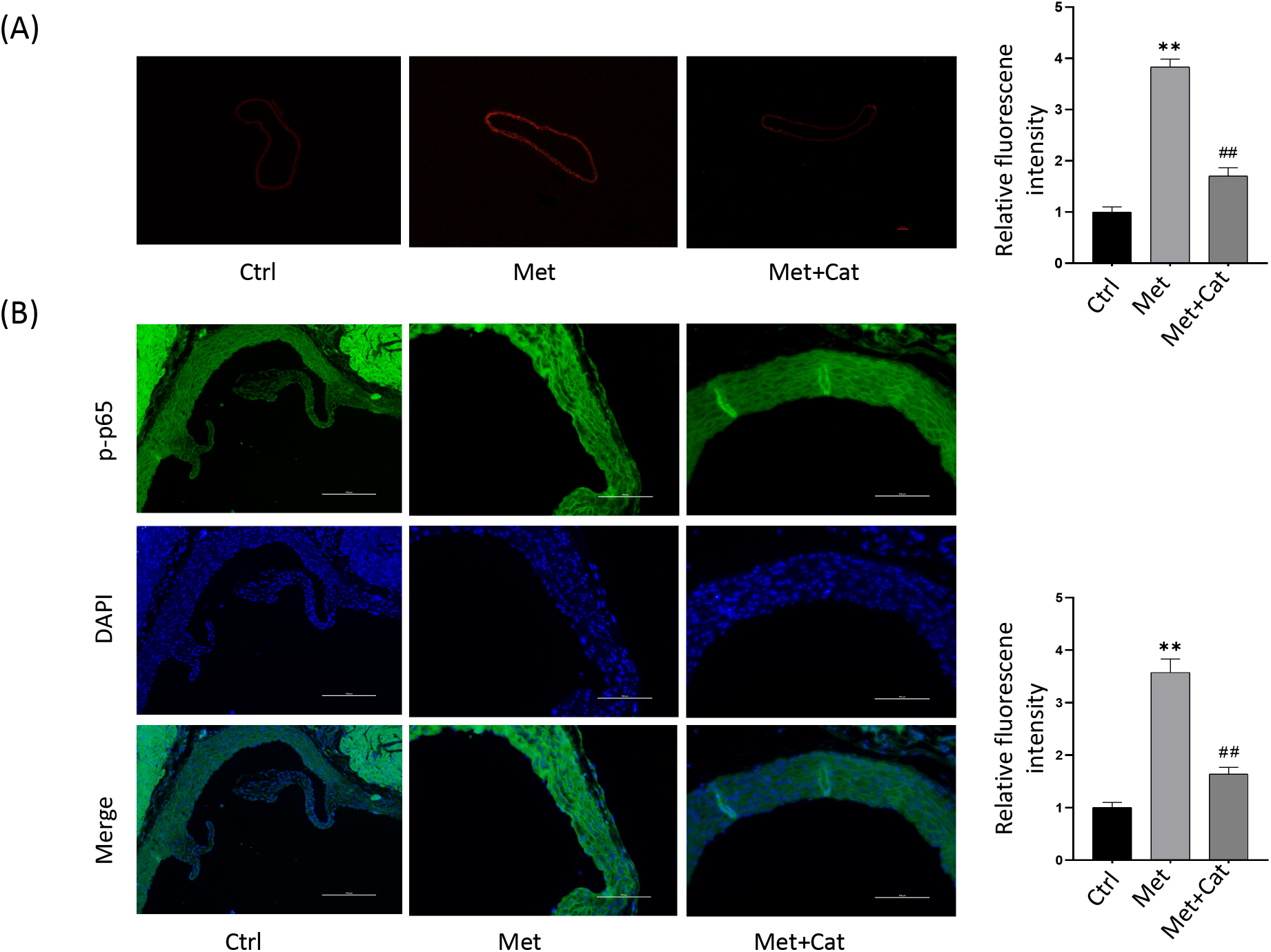
Catalpol attenuates high Met-induced EndMT by modulating ROS/NF-κB signaling. (**A**) Reactive oxygen staining of frozen tissue sections showing ROS content in aorta of control, high Met and high Met+catalpol mice, and relative fluorescence intensity (right panel). (**B**) Immunofluorescence staining showing intimal p-p65 protein expression in the aortic root of the three groups of mice, and quantification of protein fluorescence intensity (right panel). Data are expressed as mean ± SD, n = 3, *P < 0.05, ** P < 0.01, compared with the control group. #P < 0.05, ##P < 0.01, compared with the model group.

## 4 Discussion

In clinical practice, serum Hcy concentrations above 15 µmol/L are defined as HHcy, and Hcy concentrations in the ranges of 16–30, 31–100, and > 100 µmol/L are defined as mild, moderate and severe HHcy, respectively.^29^ Statistically, ∼5%–10% of the general population suffers from HHcy.^30^ The Hcy concentrations used in this study correspond to those in patients with moderate and severe HHcy in a clinical setting. The current clinical regimen for reducing HHcy levels is relatively homogeneous and mainly consists of oral folic acid and vitamin B. More than 90% of patients can reduce their Hcy concentrations within 2–6 weeks of treatment; however, the effectiveness of cardiovascular endpoint events has differed among large clinical studies.^29^ Lonn et al. and Anonymous reported that although the treatment group was able to reduce Hcy levels, it did not compare with the placebo group reductions in cardiovascular endpoint events (coronary death, myocardial infarction, or coronary regeneration).^30,31^ Therefore, although folic acid reduces Hcy levels in most patients, it is not yet clear whether it is beneficial for cardiovascular outcomes. Therefore, identification of new targets for the treatment of HHcy-related vascular diseases is essential.

EndMT is a universal feature of organ development, regeneration, and chronic fibrotic diseases, and is broadly similar to EMT in its broadest sense, which is a specific form of EMT. In recent years, there has been increasing evidence of the involvement of EndMT in the development of cardiovascular diseases, including atherosclerosis, pulmonary hypertension, valve disease, remodeling after vascular injury, and myocardial fibrosis.^20^ Evrard et al. revealed that mesenchymal cells driven by EndMT are markers of atherosclerotic lesions, found that EndMT is associated with atheromatous plaque instability, and identified EndMT as a new target for the treatment of atherosclerosis.^18^ Although a large number of studies have demonstrated that Hcy causes related vascular diseases by inducing endothelial dysfunction,^17^ few studies have suggested that EndMT may explain the pathological mechanism of Hcy-induced endothelial dysfunction.^22,23^ The results of this study confirm the involvement of EndMT in regulating HHcy-induced vascular endothelial dysfunction. It is currently believed that the activation of EndMT involves several signaling pathways, including: TGF-β signaling, oxidative stress and inflammation, cellular metabolism, non-coding RNA, epigenetics, Wnt/β-linked protein signaling, Notch signaling, and fibroblast growth factor.^21^ These pathways are mainly divided into those dependent on TGF-β signaling and those not dependent on TGF -β signaling. However, there is only sporadic evidence that Hcy upregulates TGF-β expression. For example, Sen et al. found that Hcy triggers endothelial-myofibroblast differentiation after FAK phosphorylation by inducing upregulation of TGF-β expression.^32^ However, a large body of evidence has confirmed the involvement of oxidative stress and inflammation in regulating HHcy-induced endothelial dysfunction. First, excess Hcy promotes ROS production through an autoxidation process catalyzed by metal cations such as copper.^33^ HHcy also promotes ROS production by upregulating the expression of nicotinamide adenine dinucleotide phosphate (NADPH) oxidase.^34,35^ In addition, HHcy has antioxidant activity and is associated with a decrease in enzymatic and non-enzymatic antioxidants, such as reduced expression of glutathione peroxidase, superoxide dismutase,^17,36^ glutathione, and vitamins A, C, E, and B^12^.^17^ Evrard et al. reported that endothelial cells can be converted into myogenic cells driven by oxidative stress.^18^ Collectively, these results suggest that oxidative stress may be involved in the regulation of HHcy-induced EndMT. The results of the present study confirm this hypothesis.

O_2-_ and H_2_O_2_ in ROS are well-known NF-κB agonists that promote HHcy-associated inflammatory responses. HHcy can also directly promote intranuclear migration of NF-κB family transcription factors p65 and p50 subunits to trigger inflammatory responses.^36,37,38^ In addition, HHcy has also been shown to increase the expression of inflammatory factors such as tumor necrosis factor-α, interleukin-1β upregulation, macrophage aggregation, intercellular adhesion molecule-1, monocyte chemotactic protein-1, and vascular adhesion molecule-1, which in turn promote inflammatory cell-endothelial interaction and induce a chronic inflammatory response in blood vessels.^37,38,40^ These findings suggest that HHcy impairs endothelial cell function by promoting oxidative stress and inflammation, thereby inducing vascular diseases. In the present study, ROS/NF-κB signaling was also shown to be involved in regulating HHcy-induced EndMT.

Several studies have shown that catalpol performs its biological functions by downregulating the expression of oxidative factors (MDA, PCG, LDH, AGEs, and ROS) and upregulating the expression of antioxidant factors (SOD, GSH, GSH-Px, and TAS).^41,43,44^ Catalpol was also found to inhibit P22, NOX2, and NOX4 protein expression, thereby reducing ROS production.^38,41^ Therefore, catalpol may ameliorate atherosclerosis by reducing oxidative stress. Catalpol may exert anti-inflammatory effects by downregulating the expression of inflammatory cytokines.^42,43,44,45^ In the present study, catalpol improved HHcy-induced endothelial dysfunction by restoring antioxidant levels and reducing inflammation.

In this study, we identified catalpol as a potential drug for treating Hcy-related vascular diseases. This study showed *in vitro* and *in vivo* that HHcy downregulated the endothelial markers VE-cadherin and CD31, and upregulated the mesenchymal markers N-cadherin and α-SMA, promoting increased intracellular ROS content and NF-κB nuclear translocation, which in turn induced EndMT, resulting in a morphological transition from endothelial to mesenchymal cells. While the antioxidant NAC reversed the cellular morphological transformation induced by HHcy treatment and abnormal protein expression, catalpol treatment also reversed these effects induced by HHcy. These results suggest that catalpol may exert anti-Hcy-related vascular dysfunction by attenuating HHcy-induced EndMT through the modulation of ROS/NF-κB signaling. Mechanistically, HHcy-induced oxidative stress produces large amounts of ROS, which subsequently mediate the activation of NF-κB signaling and in turn promotes nuclear transcription and induces EndMT. Interventional treatment with catalpol inhibited cell morphological transformation and the abnormal expression of EndMT-related proteins. In conclusion, catalpol protects endothelial cells from EndMT by inhibiting oxidative stress and inflammation (Figure 7b). Therefore, catalpol has a potential protective effect against HHcy-induced EndMT and endothelial dysfunction.

**Figure 7.**
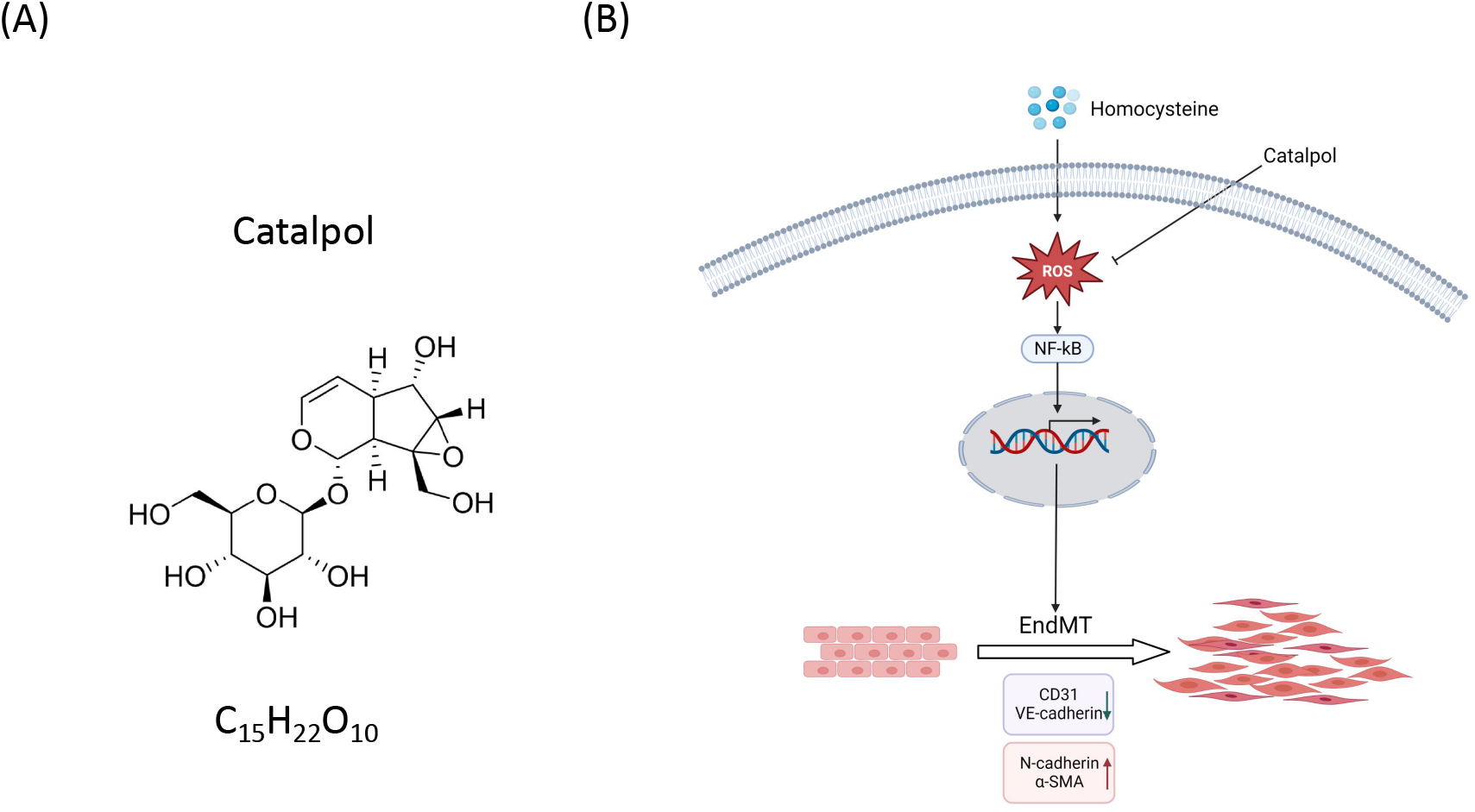
Catalpol attenuates the mechanism of HHcy-induced EndMT by inhibiting ROS/NF-κB signaling. (**A**) Chemical structure formula and molecular weight of catalpol. (**B**) Mechanism of catalpol inhibition of ROS/NF-κB signaling to attenuate HHcy-induced EndMT.

## 5 Conclusion

In summary, this study demonstrated the involvement of EndMT in the regulation of HHcy-induced endothelial dysfunction *in vitro* and *in vivo*. We also concluded that catalpol attenuated HHcy-induced EndMT by inhibiting ROS/NF-κB signaling. This study further revealed the pathological mechanism of HHcy-related vascular disease and vascular remodeling, and provided a pharmacological basis for the clinical application of catalpol.

## Author Contributions

Chengyan Wu, the first author, the experimental leader and the author of the article; Rusheng Zhao, Yuanhao Li, Peixia Li assisted in the experiments; Huibing Liu, Libo Wang, Xuehui Wang revised the article; Libo Wang, Xuehui Wang were the corresponding authors.

## Acknowledgments

We thank Xuehui Wang, Huibing Liu and Lipo Wang for their help in research and the First Affiliated Hospital of Xinxiang Medical University, Heart Center of Xinxiang Medical University for its platform.

